# Skin inflammation driven by differentiation of quiescent tissue-resident ILCs into a spectrum of pathogenic effectors

**DOI:** 10.1101/461228

**Authors:** Piotr Bielecki, Samantha J. Riesenfeld, Monika S. Kowalczyk, Maria C. Amezcua Vesely, Lina Kroehling, Parastou Yaghoubi, Danielle Dionne, Abigail Jarret, Holly R. Steach, Heather M. McGee, Caroline B. M. Porter, Paula Licona-Limon, Will Bailis, Ruaidhri P. Jackson, Nicola Gagliani, Richard M. Locksley, Aviv Regev, Richard A. Flavell

**Author notes:** These co-first authors contributed equally to this work (Piotr Bielecki, Samantha J. Riesenfeld, Monika S. Kowalczyk). Correspondence to: A.R and R.A.F.

## Abstract

Psoriasis pathology is driven by the type 3 cytokines IL-17 and Il-22, but little is understood about the dynamics that initiate alterations in tissue homeostasis. Here, we use mouse models, single-cell RNA-seq (scRNA-seq), computational inference and cell lineage mapping to show that psoriasis induction reconfigures the functionality of skin-resident ILCs to initiate disease. Tissue-resident ILCs amplified an initial IL-23 trigger and were sufficient, without circulatory ILCs, to drive pathology, indicating that ILC tissue remodeling initiates psoriasis. Skin ILCs expressed type 2 cytokines IL-5 and IL-13 in steady state, but were epigenetically poised to become ILC3-like cells. ScRNA-seq profiles of ILCs from psoriatic and naïve skin of wild type (WT) and *Rag1*^-/-^ mice form a dense continuum, consistent with this model of fluid ILC states. We inferred biological “topics” underlying these states and their relative importance in each cell with a generative model of latent Dirichlet allocation, showing that ILCs from untreated skin span a spectrum of states, including a naïve/quiescent-like state and one expressing the *Cd74* and *Il13* but little *Il5*. Upon disease induction, this spectrum shifts, giving rise to a greater proportion of classical *Il5-* and *Il13-* expressing “ILC2s” and a new, mixed ILC2/ILC3-like subset, expressing *Il13, Il17,* and *Il22*. Using these key topics, we related the cells through transitions, revealing a quiescence-ILC2-ILC3s state trajectory. We demonstrated this plasticity *in vivo*, combining an IL-5 fate mouse with IL-17A and IL-22 reporters, validating the transition of IL-5–producing ILC2s to IL-22– and IL-17A–producing cells during disease initiation. Thus, steady-state skin ILCs are actively repressed and cued for a plastic, type 2 response, which, upon induction, morphs into a type 3 response that drives psoriasis. This suggests a general model where specific immune activities are primed in healthy tissue, dynamically adapt to provocations, and left unchecked, drive pathological remodeling.

## INTRODUCTION

Psoriasis pathology is driven by the type 3 cytokines IL-17 and Il-22 (Nograles, Zaba et al. 2008, Cai, Shen et al. 2011). The dermal inflammation and acanthosis induced by IL-23 are thought to be mediated by the Th17-cell associated cytokine IL-22. While both IL-17 and IL-22 are produced at elevated levels in psoriatic skin, the major cell source of these cytokines during disease remains unclear. In psoriasis patients, γδ T cells were found to be greatly increased in affected skin and produced large amounts of IL-17, suggesting they may play a key role in pathogenesis (Cai, Shen et al. 2011). Recently, however, IL-17 and IL-22 producing ILC3s have been proposed to be a significant source of cytokine production during psoriasis (Teunissen, Munneke et al. 2014, Villanova, Flutter et al. 2014). These cells are thought to be activated in response to IL-1β and IL-23, whose levels correlate with disease severity, and are decreased following antitumor necrosis factor-α (anti-TNFα) treatment. The presence of a novel ILC population in psoriatic skin that responds to one of the most effective biologic therapeutics suggests that dysregulation of ILCs is a contributing factor to psoriasis pathogenesis. While ILC3s dominate psoriatic skin (Pantelyushin, Haak et al. 2012), in healthy individuals the majority of ILCs are represented by group 2 ILCs, defined by IL-5 and IL-13 production (Roediger, Kyle et al. 2013, Spencer, Wilhelm et al. 2014). We wanted to determine at what point during disease progression the frequency of ILC2s and ILC3s shifts and whether a potential for conversion between these cell types underlies this disease.

High parallel single-cell RNA sequencing (scRNA-seq) has become a powerful tool for unbiased analysis of various cells types. Analyses of immune cell classes, such as ILCs, typically treat them as collections of discrete immune cell “types”, yet these cell types may share important biological signals and have been observed in some contexts to essentially continuously span a functional spectrum. To capture and explore fluid, mixed transcriptional states, we used latent Dirichlet allocation (LDA), or “topic modeling”, a statistical data mining approach for discovering the abstract topics that explain the words occurring in a collection of text documents (Blei 2003). Applied to scRNA-Seq, each “document” corresponds to a cell, and a “topic” corresponds to a biological program, modeled as a distribution over expressed genes, rather than words. Given the number of topics as a parameter, both topics and the mixture weights in cells are inferred without supervision. LDA was independently introduced in population genetics to model admixed individuals with ancestry from multiple populations (Pritchard, Stephens et al. 2000). In genomics, it has been applied to deconvolute cell types in population RNA-seq (Repsilber, Kern et al. 2010, Schwartz and Shackney 2010, Shen-Orr, Tibshirani et al. 2010, Ahn, Yuan et al. 2013, Lindsay 2013, Quon, Haider et al. 2013, Wang, Gong et al. 2015), and proposed for finding structure in bulk or single-cell RNA-seq, for example, in inference of confounding batch effects (Dey, Hsiao et al. 2017).

## RESULTS

To determine which cells are key to initiate psoriatic disease, we studied a subcutaneous IL-23 injection model, which leads to increased skin thickness after five days of daily injections (**Fig. 1A**, **Fig. S1A**). First, we assessed the role of different immune cell types in this model (**Fig. 1B**, **Fig. S1B and C**). Consistent with previous results, the *Rag2*^-/-^ *Il2rg*^-/-^ double mutant, which lacks all lymphocytes, did not show any increase in ear thickness, whereas *Rag1*^-/-^ mice, which have intact ILCs, showed significant increase in skin thickness over the treatment course. This is also consistent with an increased number of human ILC3s recently observed in psoriatic patients (Teunissen, Munneke et al. 2014, Villanova, Flutter et al. 2014). Moreover, while γδ T cells have been implicated in a longer treatment course (Cai, Shen et al. 2011, Pantelyushin, Haak et al. 2012), analyzing *Tcrd*^-/-^ mice, which lack only γδ T cells, we found no evidence that they contribute to disease initiation (**Fig. 1B**). Next, to further confirm the role of ILCs in disease initiation, we adoptively transferred sorted skin ILCs from untreated WT mice into *Rag2*^-/-^ *Il2rg*^-/-^ mice, and observed significant skin thickening in treated versus untreated recipient mice (**Fig. 1C**). Finally, we assessed the contributions of circulatory versus tissue-resident lymphocytes in the psoriasis model, because recent studies of inflammation in several peripheral tissues suggested different involvement of circulatory and tissue resisdent ILCs (Dyring-Andersen, Geisler et al. 2014, Gasteiger, Fan et al. 2015, Li, Hodgkinson et al. 2016, Yang, Hu et al. 2016, Huang, Mao et al. 2018). We compared disease phenotype between control mice and those treated with FTY720, which blocks signaling from the S1P1 receptor, preventing egress of T cells from secondary lymphoid tissues and limit trafficking of induced ILC2s (Matloubian, Lo et al. 2004, Huang, Mao et al. 2018). FTY720-treated mice had the expected reduction of circulating total white blood cells, but showed no difference in psoriasis phenotype induction upon IL-23 administration compared with untreated controls, in both WT or *Rag1*^-/-^ (lacking T and B cells) mice (**Fig. 1C and E**). Thus, in contrast to a model of lung inflammation (Huang, Mao et al. 2018), in psoriasis, tissue-resident ILCs are sufficient to drive disease pathology and are critical for amplifying the response to IL-23.

**Figure 1.**
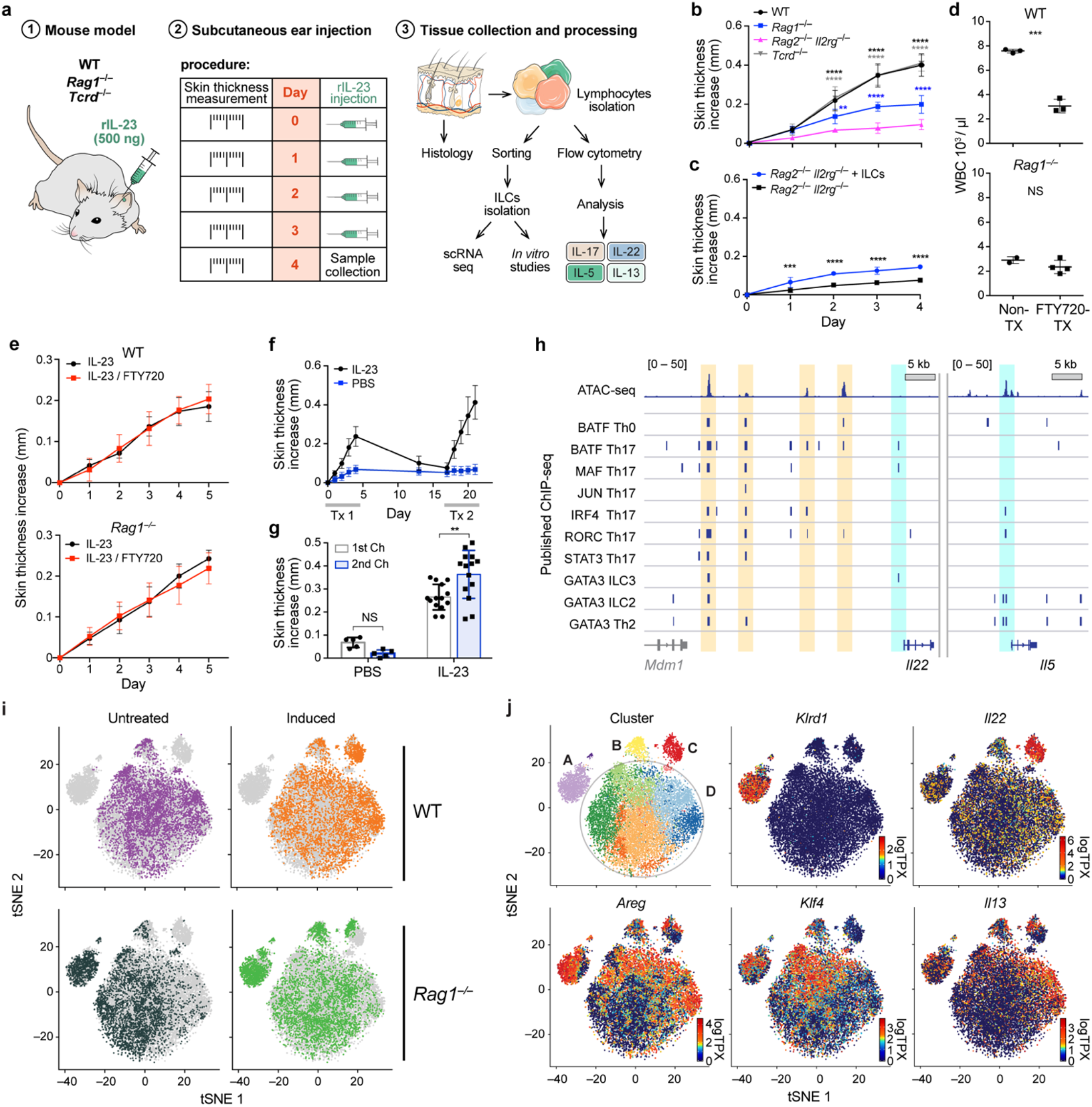
An epigenetically poised, heterogenous population of tissue-resident ILCs drive initial IL-23–induced pathology. **(A)** Study overview. From left: Psoriasis mouse model is based on a series of subcutaneous IL-23 injections in WT, *Rag1*^-/-^ and *Tcdr*^-/-^ mice, phenotypic measurement of skin thickness, and tissue collection and cell isolation for assessment by scRNA-seq, in vitro assays and cytokine expression. (**B,C)** Tissue resident ILCs are necessary and sufficient for increase in ear skin thickness in response to IL-23 treatment. Increase in skin thickness (mm, y axis) over time (days, x axis) in (**B**) WT (black), *Rag1*^-/-^ (lack all T and B cells, blue), *Rag2*^-/-^ *Il2rg*^-/-^ mice (also lack ILCs, magenta), and TCRγδ^-/-^ (lack γδ T cells, grey) (n=7 for each group) as well as in (**C**) *Rag2*^-/-^ *Il2rg*^-/-^ mice with (blue) and without (black) intravenously transferred ILCs (n=4 for each group)**. (D)** FTY720 blocks white blood cell circulation. Total circulatory white blood cell (WBC) numbers (10^3^/μl; y axis) in untreated (“Non-Tx”) and FTY720-treated (“FTY720-Tx”), WT and Rag^-/-^ mice. (**E)** IL23-dependent increases in ear skin thickness does not require circulating cells. Increase in skin thickness (mm, y axis) over time (days, x axis) following IL-23 treatment, in WT (top) and Rag^-/-^ (bottom) mice, with (red) and without (black) FTY720-treatment (**Methods)** (n=3 WT both groups, n=2 *Rag1*^-/-^ NonTX n=4 *Rag1*^-/-^ FTY720). (**F,G**) A secondary challenge with IL-23 increases susceptibility. Increase in skin thickness (mm, y axis) over time (days, x axis; top) or at the end (bottom) of a primary (white bar) or secondary (blue bars) challenge with either IL-23 (n=14) or saline control (PBS) (n=5). (**H)** ILCs in untreated mice are epigenetically poised to become ILC3s. Mapped ATAC-seq reads (top track) at the *Il22* (left) and *Il5* (right) promoter loci (bottom track) from sorted skin ILCs from untreated mice, show open chromatin peaks (beige and blue bars) at key TF binding sites (beige), previously identified in CD4^+^ T cell ChIP-seq data (middle tracks), and at the TSS of *Il5* but not *Il22* (blue). (**I,J)** ILC heterogeneity highlighted by scRNA-seq. *t*-Distributed stochastic neighbor embedding (tSNE) of 27,998 single cell (dots) profiles (**Methods**) colored by either *in vivo* treatment and genotype (**i**), or by cluster assignment or expression of key genes (color bar, logTPX (**Methods**)) (**J**). Annotated clusters (**J**, top left) include a *Rag1*^-/-^-specific cluster (A) expressing the ILC1-associated gene *Klrd1*, cycling cells (B), an *Il22-*high cluster co-expressing *Il13* (C), and a heterogeneous “cloud” (D), without discrete boundaries between clusters yet with multiple patterns of graded gene expression. Error bars, SD; **p<0.021, ***p<0.0002, ****p<0.0001 by unpaired t test (b) or two-way ANOVA (d, e, g).

We observed that skin ILCs expressed the type 2 cytokines IL-5 and IL-13 in steady state, but showed potential to plastically to assume ILC3-like states. Consistent with prior reports that naïve mouse skin ILCs are comprised almost exclusively of GATA3+ ILC2s (Roediger, Kyle et al. 2013), total ILCs isolated from healthy mouse skin and treated with the type 2 alarmin cytokines IL-25 and L-33 had a strong type 2 activation, as indicated by expression of *Areg* and *Il13* (**Fig. S1D**). However, total ILCs treated with IL-23 and IL-1β instead strongly expressed *Il22* and *Il17a* (**Fig. S1D**), suggesting that tissue-resident skin ILCs may have potential for type 2-3 plasticity. Such plasticity has been previously reported in IL-17A co-expressing “inflammatory ILC2s” in the lung (Huang, Guo et al. 2015, Zhang, Xu et al. 2017), similar to reported type 3-1 plasticity in gut and tonsil ILCs and type 2-1 plasticity in blood ILCs (Cella, Otero et al. 2010, Bernink, Krabbendam et al. 2015, Bal, Bernink et al. 2016, Lim, Menegatti et al. 2016, Ohne, Silver et al. 2016, Silver, Kearley et al. 2016). Moreover, while inflammation and skin thickness reverted to near-baseline levels within 10 days after the initial IL-23 injection (**Fig. 1F**), this initial challenge promoted a stronger type 3 response upon re-challenge. Specifically, mice showed a significantly more severe phenotype after a second series of IL-23 injections, compared to their initial response (**Fig. 1F and G**). This was also observed in mice treated with FTY720 during the primary injection (**Fig. S1E**), suggesting that the plastic psoriatic response is not due to ILC recruitment.

We hypothesized that this plasticity may be encoded epigenetically. To test this hypothesis, we profiled sorted total skin ILC populations from naïve mice by ATAC-seq. We observed the expected open chromatin signature at the TSS of *Gata3*, *Il5* and *Il13* and not at the TSS of *Tbx21* or *Rorc,* which encode T-bet and Rorγt, the hallmark transcription factors (TFs) of ILC1s and ILC3s, respectively, or at the TSS of *Il22*, *Il17a*, or *Il17f* (**Fig. S1F and G**). In support of our hypothesis, we also observed strong ATAC-seq peaks at at promoters of some type 3 genes in TFs binding sites, such as *Batf, Maf,* and *Irf* (Ciofani, Madar et al. 2012, Li, Spolski et al. 2012, Zhong, Cui et al. 2016) (**Fig. 1H**, **Fig. S1G**), which are known to regulate Th17 cells. Taken together, our data support a model where IL-23 induces psoriasis by remodeling a heterogeneous, tissue-resident ILC population with unexpected potential for differentiation, rather than by recruiting circulating ILCs to replace a homogenous, terminally differentiated skin-resident ILC2 population.

To assess the molecular heterogeneity of skin-resident ILCs and its functional implications for the IL-23 response, we collected massively parallel scRNA-seq profiles from sorted pure total ILCs from WT and *Rag1*^-/-^ mice from naïve and IL-23 induced conditions, predominantly uncovering a large heterogeneous population of cells (**Fig. 1I**). Specifically, clustering on principle components, followed by differential expression analysis (**Methods**), identified a few discrete subsets of cells, including a *Rag1*^-/-^-specific subset (*A*), a cluster of proliferating cells from all conditions and genotypes (*B*), and a cluster specific to the induced condition with very high *Il22* expression and some *Il13* expression (*C*) (**Fig. 1J, Fig. S1H**). However, the vast majority of cells (81%) formed a single, large heterogeneous and continuous “cloud” (*D*), which was not simply driven by technical factors (**Methods**), with multiple sub-regions enriched for specific functional programs, including type 2 immune response (**Fig. 1J**). Importantly, no single partitioning conformed to the expression of key genes and processes, and moreover, some biological processes were unexpectedly shared across subsets of the cells from distinct clusters (**Fig. 1J**). This highlighted the diversity of potential cell states, and the need to capture them by more nuanced computational analysis.

To characterize the heterogeneity of ILCs during IL-23 response, we created a generative topic model based on LDA. Analogous to a text document, a cell is modeled as a mixture of a small number of topics, where the mixture weights indicate the relative prominence of the corresponding biological process in that cell. Multiple topics may include the same gene, reflecting the gene’s roles in different processes. Topic modeling permits a cell to have multiple, non-hierarchical “identities” that potentially differ in importance, a feature particularly relevant for analyzing cellular plasticity (**Fig. 2A**). Indeed, we observed complex patterns of topic sharing across clusters, suggesting that topic weights capture relationships not well described by clusters and, through their functional interpretation, enable a more nuanced view of similarities and differences among cells (**Fig. 2B**). Several choices for the number of topics may result in valid models, though too large a number of topics can result in overfitting and low interpretability. We found that in this dataset, 15 topics captured important changes during disease induction, as well as other signals, without obvious signs of overfitting (**Fig. S2A**, **Methods**).

**Figure 2.**
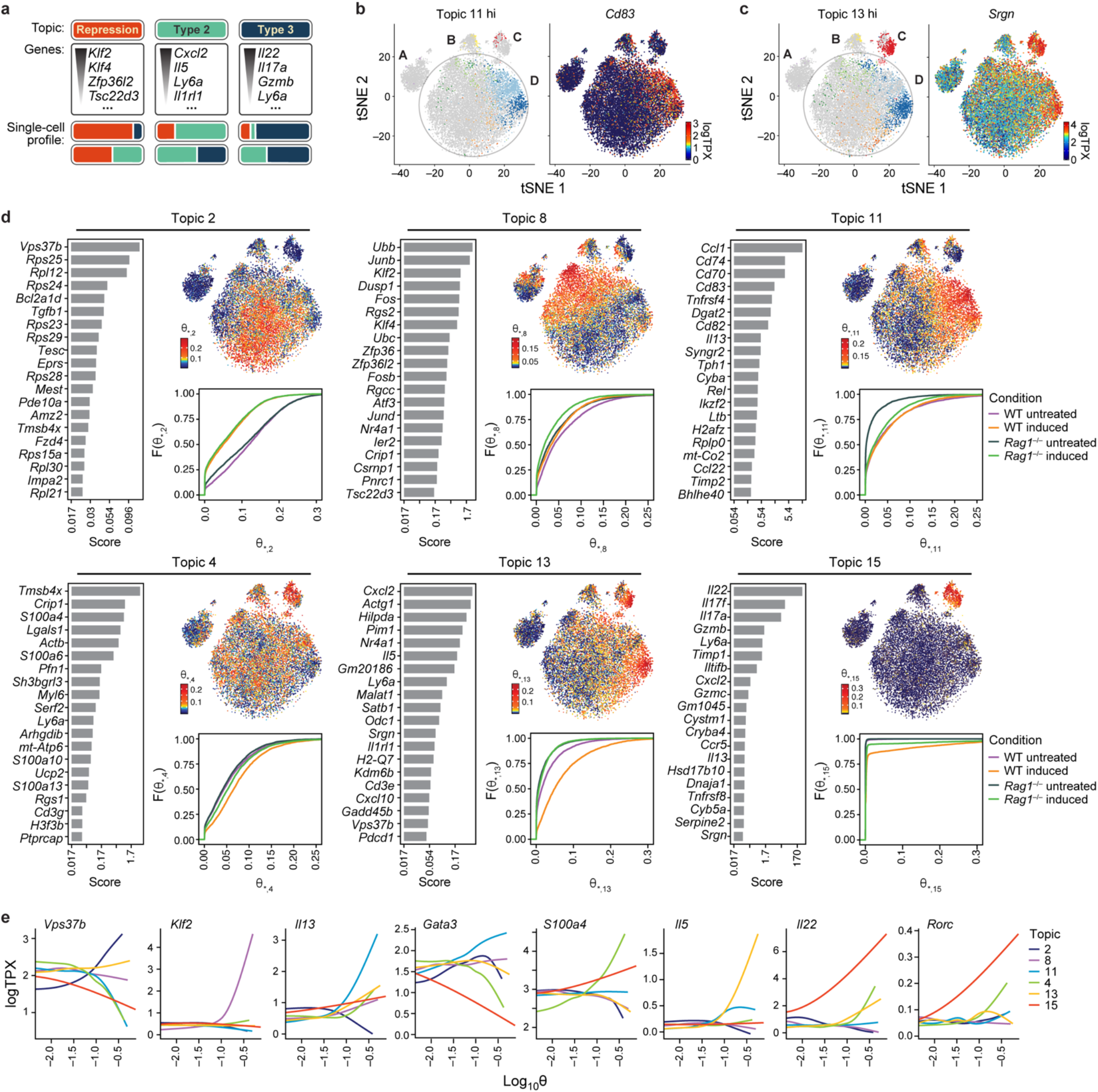
Topic modeling of skin ILCs highlights repressive, quiescent-like state and multiple, distinct states of activation combined in cells. **(A)** Topic model concept in the context of single cell expression. Topics (top) consist of genes (middle), with distinct weights (gradient, **Methods**) based on their importance in the topic. Cells (bottom), are scored based on the contribution of each topic (color) in them; a cell can thus have multiple topics. (**B–E),** Results of LDA on ILCs with 15 topics (**Methods**). (B,C) Topics reveal complex relationships among clusters. TSNE of cells colored if they are highly weighted for Topic 11 and gray otherwise, with color code reflecting cluster membership as in **Fig. 1J** (**B**, left), or by expression (color bar, logTPX (**Methods**)) of *Cd83*, a Topic-11 associated gene (**b,** right). Analogous plots for Topic 13 (**c,** left) and its associated gene *Srgn* (**C**, right). (D,E) Topics with high weights in cells from untreated (Topics 2, 8, and 11) *vs*. induced (Topics 4, 13, and 15) conditions. (**D**) For each topic shown are a bar plot of top scoring genes (y axis), ranked by a score (x axis, logarithmic scale) of how well the gene distinguishes this from other topics (**Methods**); a tSNE (as in **Fig. 1I**) with cells colored by the topic’s weight in the cells (column *j* of the cell-by-topic weight matrix θ (θ_*_,*j*) for Topic *j*); and a graph of the empirical cumulative density function (y axis) of topic weights θ_*_,*j* (x axis) for cells grouped by treatment or genotype (as in **Fig. 1H**). (**E**) Examples of topic-associated genes. Gene expression (y axis, logTPX) as a smoothed function of the topic weight (log θ_*_,*j*, x axis), for each of the topics highlighted in **D** (color)

Our topics spanned three categories: (**1**) highly ribosomal- or mitochondrial-dominated (*e.g.*, Topic 1, 6), possibly reflecting technical quality or cell size, (**2**) cluster-specific topics (*e.g.*, Topic 7, 14, 15), and (**3**) “sub-regional” topics, that is, those featured in sub-regions of the “cloud”, also often simultaneously present in sub-regions of other clusters (*e.g.*, Topics 2, 4, 8, 11, 13) (**Fig. 2C and D**, **Fig. S2B–D**). “Cell quality” topics can help distinguish the influence of technical confounders better than simple thresholds, but also may reflect a cell’s level of biological activation (Wallrapp, Riesenfeld et al. 2017). “Cluster-specific topics” are analogous to results from standard differential expression analysis. For example, cluster *C* is unique in having large weights for Topic 15, which is characterized by expression of ILC3-associated genes *Il22, Il17a,* and *Il17f,* as well as the cytotoxic gene *Gzmb* and the type 2 genes *Ly6a* (Sca-1) and *Il13* (**Fig. 2C and D**, **Fig. S2B and C**). As another example, Topic 7 is uniquely highly weighted in cells from the *Rag1*^-/-^-specific cluster *A*, and features the NK-associated genes *Klrd1* and *Tyrobp* and the immunoglobulin E receptor *Fcer1g,* indicating that *Rag1*^-/-^ mice might have an overrepresentation of skin-resident ILC1s (**Fig. 1J**, **Fig. S2D**).

The “sub-regional” topics highlighted functional states that are prominent within the “cloud” and span across cluster boundaries, showing that ILCs from untreated skin span a spectrum of immune states, including one characterized by *Vps37b* expression (Topic 2), a naïve/quiescent-like state (Topic 8) and an activated state related to antigen presentation (Topic 11). Notably, this may mirror "functional compartmentalization" reported in gut ILCs in homeostasis (Gury-BenAri, Thaiss et al. 2016). This spectrum shifted upon disease induction, giving rise to greater representation of classical *Il5*- and *Il13*-expressing “ILC2s” (Topic 13), as well as a mixed ILC2/ILC3-like state characterized by strong expression of *Il13*, *Il17*, and *Il22* (Topic 15) (**Fig. 2C and D, Fig. S2B**). Specifically, Topic 2, mainly present in the “cloud”, distinguishes between the untreated and induced conditions, partly through ribosomal genes that may reflect differences in size between naïve and activated cells (**Fig. 2C**). Topic 8 is characterized by expression of TFs previously associated with both T- or B-cell quiescence, such as *Klf2/Klf4* (Carlson, Endrizzi et al. 2006, Cao, Sun et al. 2010) and *Zfp36l2* (Galloway, Saveliev et al. 2016, Salerno, Engels et al. 2018), and with repression of Th17 genetic programs, such as *Tsc22d3* (Yosef, Shalek et al. 2013), and may thus reflect an actively maintained quiescent ILC state (ILC0) (**Fig. 2C and D**, **Fig. S2C**). Topic 11, which is present in cells from both WT conditions and the *Rag1*^-/-^ induced condition, features genes associated with antigen presentation, including MHCII invariant chain and MIF receptor *Cd74(Schroder 2016)* and *Cd83* (Kuwano, Prazma et al. 2007), and type 2 ILCs (e.g., *Il13, Ccl1,* and *Dgat2*, though not *Il5*) (Robinette, Fuchs et al. 2015, Gury-BenAri, Thaiss et al. 2016, Wallrapp, Riesenfeld et al. 2017, Ricardo-Gonzalez, Van Dyken et al. 2018) (**Fig. 2C and D**). Topic 13, highlighting a substantial sub-region of both the “cloud” and induced-specific cluster *C*, is more specific to WT disease induction, uniquely expresses *Il5*, and also includes other type 2 genes, such as *Cxcl2*, *Il1rl1* (ST-2), *Il13*, and *Ly6a* (Sca-1), the latter of which featured in all induced topics (**Fig. 2C and D**, **Fig. S2B and C**). The presence of some cells with high weights for both Topics 13 and 15 indicates that an activated type 2 response apparently co-exists with the anticipated type 3 response. Finally, Topic 4, which is largely mutually exclusive with Topic 13 across cells, includes genes involved in actin remodeling, a process previously shown to be important during T-cell activation (Kumari, Curado et al. 2014) (**Fig. 2C and D**).

We hypothesized that cells can transition between some of these programs or states, as such transitions would be consistent with the dense transcriptional continuum observed. Unlike pseudotime inference (Trapnell, Cacchiarelli et al. 2014, Haghverdi, Buttner et al. 2016), topic modeling does not assume the existence of an “axis" of progression, which may not exist in settings such as the untreated condition. Moreover, when a trajectory does exist, it may be reflected only in specific aspects of the transcriptional profiles. Indeed, a temporal “induction” dimension in our data was revealed most clearly when we focused on specific topics related to immune repression or activation. To identify transitional relationships in the context of the biological processes reflected by these topics, we created a diffusion map only from those cells highly weighted for Topics 2,4,8,11,13, and 15, but not for Topics 6 or 7, and used only the most distinguishing genes for each topic as input (**Fig. S3A-C**, **Methods**).

The diffusion map (**Fig. 3A, Fig. S3D**) proposes several parallel state transitions that cells undergo in the tissue, in particular highlighting a quiescence-ILC2-ILC3s state trajectory in the disease. **First**, cells from the naïve condition lie in a triangular region in the plane spanned by diffusion components (DC) 2 and 3 with corners up-weighted for Topic 2 (“resting”), 8 (“naïve-quiescent”), and 11 (“antigen presentation”), respectively (**Fig. 3A and B.i–iii**). Their distribution throughout the triangle suggests that in the untreated condition, cells range over all mixtures of these states. **Second**, DC1 captures the induced response shared in both WT and *Rag1*^-/-^ mice (**Fig. 3A, Fig. S3D**), such that as their DC1 coordinate (“induction”) increases, cells typically have relatively lower weights for Topics 2, 8, and 11 (**Fig. 3B.i–iii, Fig. S3E.i-iii),** and higher weights for Topic 15 (“*Il22/Il117*”), Topic 4 (“actin remodeling”), and, specifically for cells from WT mice, Topic 13 (“*Il5/Cxcl2*”) (**Fig. 3B.iv-vi**, **Fig. S3E.iv-vi**). Genotype-specific differences in the induction response are further captured by DC4, such that cells from WT and *Rag1*-/-mice have increasingly different DC4 coordinates as DC1 coordinate increases (**Fig. S3D**).

**Figure 3.**
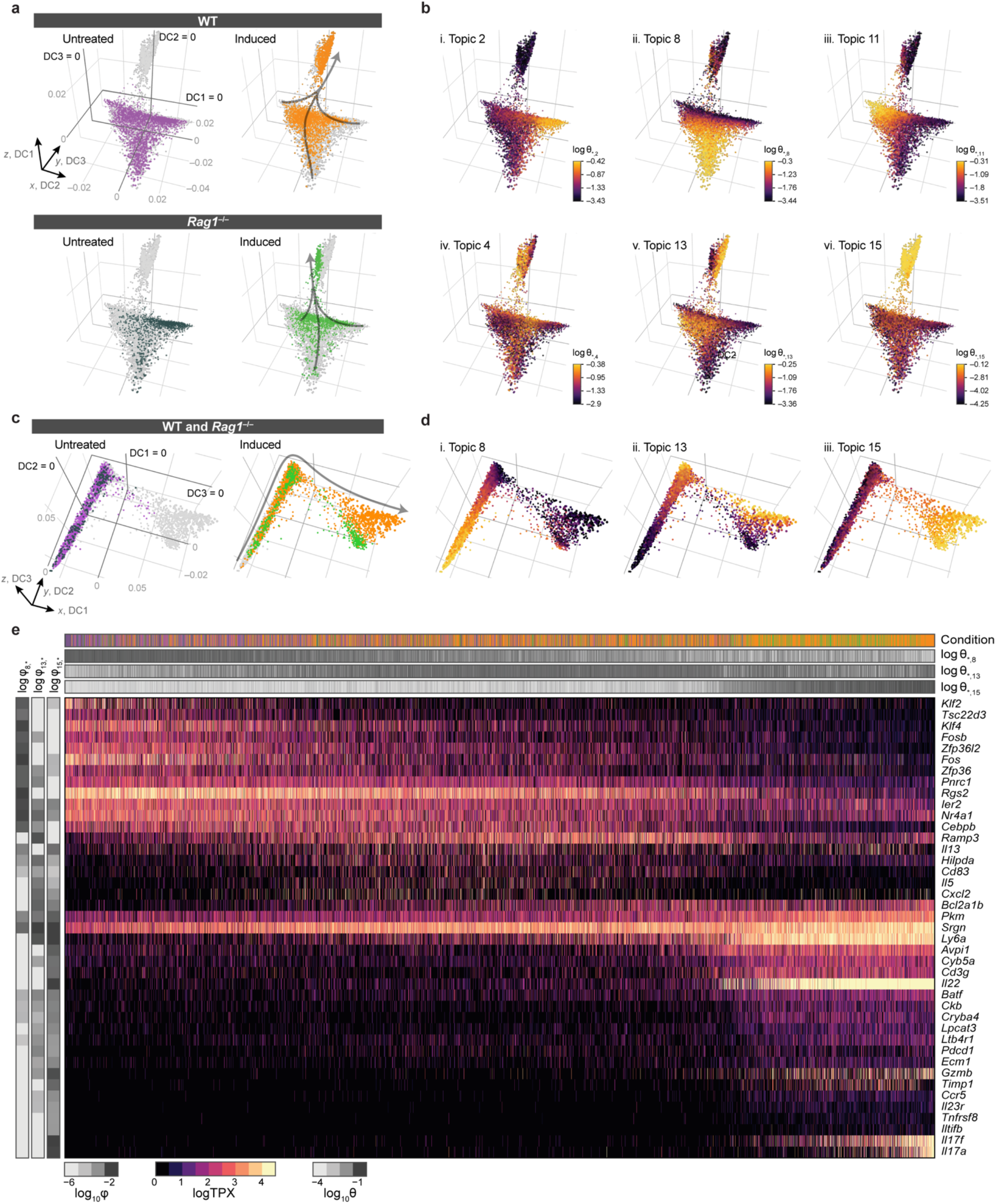
Inference of an IL-23–induced dynamic trajectory from quiescent-like ILCs through classically activated ILC2s to pathological *I13/Il17a/Il22-*expressing ILC3-like cells. **(A,B)** Distinct topics suggest a dense continuum of states undergoes a dynamic transition during psoriasis induction. Shown is a diffusion map constructed only from cells highly weighted for selected topics (Topics 2, 4, 8, 11, 13, or 15) and the corresponding topic-specific genes (**Methods**). Plots of DC2 (x axis), DC3 (y axis) and DC1 (z axis), show cells (dots) colored by either *in vivo* treatment and genotype (**A**) or by topic weight (log θ_*_,*j*, color bar) (**B**). Gray arrows (**A**) indicate an implicit direction of induction. **(C,D)** A naïve-induced trajectory across DC1 in a focused diffusion map from Topics 8, 13, and 15. DC 1 (x axis), DC2 (y axis), and DC3 (z axis) of a focused diffusion map, with cells colored as in **A** by *in vivo* treatment and genotype (**C**), or as in **B** by topic (**D**). **(E)**, Key genes associated with the trajectory from quiescent-like ILCs to activated ILC2s to ILC3-like cells. Expression (color scale, logTPX) of genes (rows) in cells (columns) associated with Topics 8 (“naïve-quiescent”), 13 (“*Il5*/*Cxcl2*”), and 15 (“*Il22/Il17a*”), with cells marked by *in vivo* condition and genotype (top bar; colored as in **A**). Grey scale bars: Topic weights for cells (log θ_*_,*j*) (horizontal bars) and genes (log β_*j*_,*, where β is the topic-by-gene weight matrix; vertical bars) illustrate mixtures of functional states.

A focused diffusion map model (**Fig. 3C**) generated only from cells up-weighted for Topic 8, 13, or 15 (**Methods**), shows continuous expression changes from Topic 8 to 13 to 15, as DC 1 (in this map) coordinate increases (**Fig. 3D and E**). Indeed, DC1 is particularly well correlated with expression of the gene *Srgn*, a proteoglycan that is critical for the trafficking and storage of *Gzmb* (Sutton, Brennan et al. 2016), which suggests that expression of this gene could be an early indicator of a trajectory toward type 3 activation, visible before expression of either *Gzmb* or type 3 cytokines (**Fig. 3E**). The expression changes observed across Topic 8, 13, and 15 are consistent with a novel model of immune activation in which a type 3 stimulus (IL-23) causes skin-resident naïve/quiescent ILCs to undergo type 2 activation, followed by transition to ILC3-like cells.

Finally, we tested the model’s predictions of a quiescent-ILC2-ILC3 trajectory. First, we validated the quiescent state by ATAC-Seq of sorted total skin ILC populations from naïve mice. Consistent with Topic 8 (“naïve-quiescent”) highlighted by the scRNA-seq analysis, the loci for the TFs *Klf2, Klf4*, previously associated with quiescence (Carlson, Endrizzi et al. 2006, Cao, Sun et al. 2010)*, Tsc22d3* and *Zfp36l2,* associated with Th17 genetic program repression (Yosef, Shalek et al. 2013, Galloway, Saveliev et al. 2016, Salerno, Engels et al. 2018) and *Cebpb*, involved in hematopoiesis (Tsukada, Yoshida et al. 2011), had open chromatin signatures at their TSS (**Fig. 4A**).

**Figure 4.**
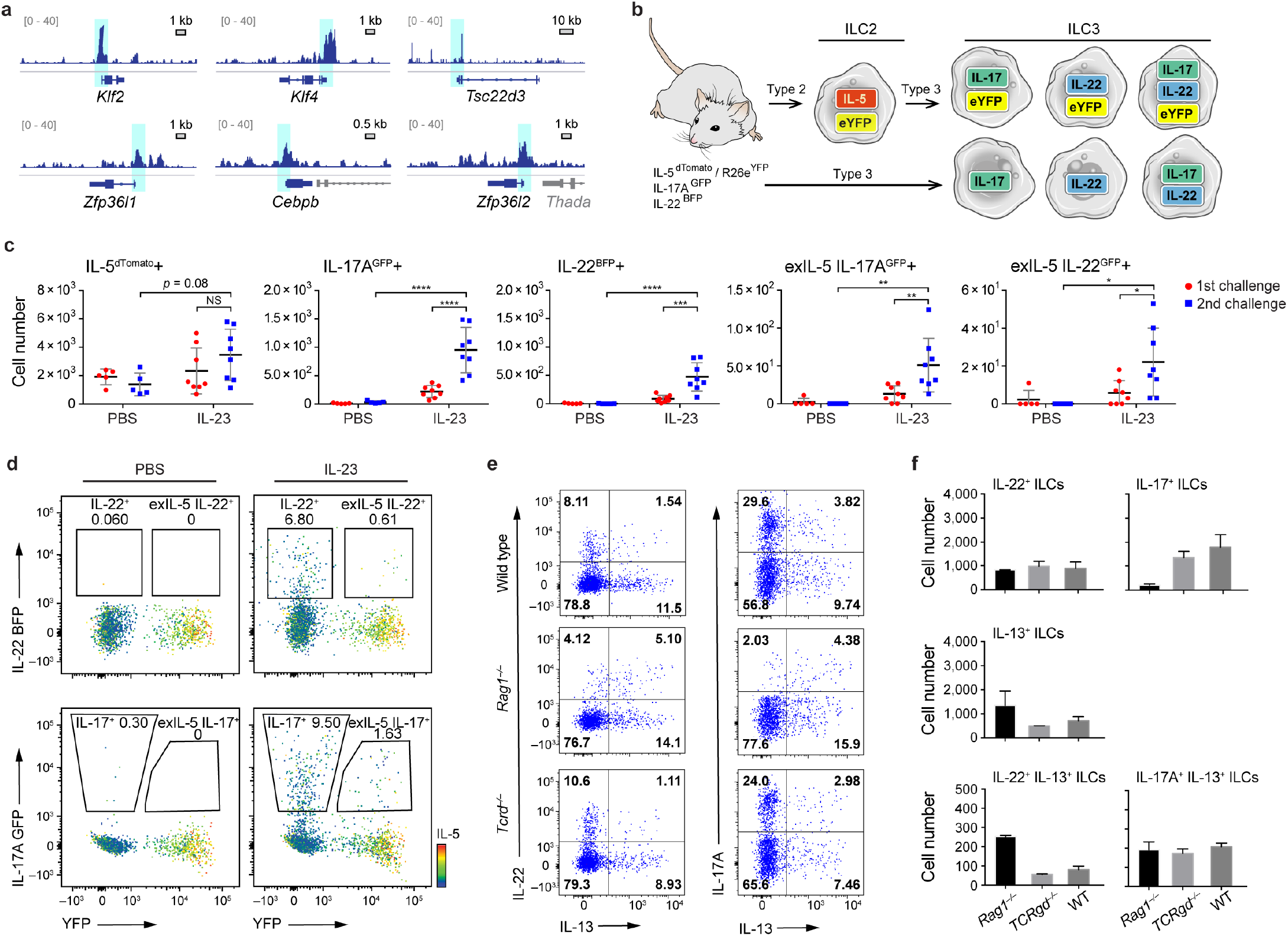
*In vivo* validation of the trajectory from quiescent-like ILCs in healthy skin to differentiation of ILC2s to ILC3-like cells during IL-23–induced response. **(A)** ATAC-seq confirms quiescent-like ILCs. Open chromatin peaks (light blue shaded boxes) of ATAC-Seq reads (blue tracks) from sorted skin ILCs from untreated mice at TSS of key genes (bottom track) responsible for quiescence and repression of type 3 programs. **(B-D)** ILC2-ILC3 plasticity revealed by IL-5 fate mapping and IL-22^BFP^ and IL-17A^GFP^ reporter mouse. (**B**) Fate mapping scheme. IL-5 Fate mouse reporter combined with IL-17A^GFP^ and IL-22^BFP^ reporters showing possible outcomes of skin ILC activation, in a scenario with ILC2 to ILC3 differentiation (top) *vs*. direct ILC3 differentiation (bottom). (**C**) IL-23 induction increases the number of IL-22– and IL-17A–producing cells, including among cells formerly producing IL-5 (“exIL-5”), especially after secondary challenge. Number of cells (y axis) with each reporter configuration (top label) in IL23-treated and PBS controls (x axis) in the first (circles) and second (squares) challenge. **(D)** exIL-5 cells that transdifferentiated to produce IL-22 and IL-17A do not produce IL-5 anymore. FACS plots of the expression of YFP (x axis) and IL22-BFP (x axis, top) or IL17A-GFP (x axis, bottom). (**E,F),** IL-23 treatment induces IL-13/IL-22 and IL-13/IL-17A double-producing populations and elevates IL-13/IL-22 double production in *Rag1* deficient mouse. (**E**) Levels of IL-13 (x axis) and IL-22 (y axis) measured by intracellular cytokine staining of skin ILCs in wild type (top), *Rag1^-/-^* (middle) and *Tcrd^-/-^* (bottom) mice. (**F**) Mean number of cells (y axis) among single producers and co-producers in each mouse genotype (x axis). Error bars, SD; *p<0.0332, **p<0.021, ***p<0.0002, ****p<0.0001 by two-way ANOVA.

Next, we tested the prediction of a transition during disease of IL-5–expressing ILC2s into IL-22/IL-17A–expressing ILC3-like cells. We generated an IL-5 fate reporter mouse from IL-5-cre-dTomato (Red5) (Nussbaum, Van Dyken et al. 2013) and Rosa26^flox-Stop-floxYFP^, which we then combined with IL-17A^GFP^ (Esplugues, Huber et al. 2011) and IL-22^BFP^ expression reporters (**Fig. 4B**). Consistent with our model, after IL-23 injection, ~10% of the IL-22– and IL-17A–expressing cells were indeed ex-IL-5 producing cells, as measured by fate mapping of ILC2s, and a second IL-23 challenge further elevated the number of ex-IL-5 cells producing IL-22 and IL-17A (**Fig. 4C and D**). Moreover, cells that expressed ILC3 type cytokines no longer expressed IL-5 (**Fig. 4D**). Our results show the *in vivo* potential for plasticity among skin ILCs and demonstrates that some cells expressing ILC3 type cytokines expressed IL-5 at one stage of their lifetime. Finally, we also tested our model’s prediction that there is a subset of skin ILCs in the psoriasis model that co-expresses the type 2 cytokine IL-13 with both of the type 3 cytokines IL-22 and IL-17A. Indeed, intracellular measurements of these three cytokines showed that, consistent with the predictions, nearly 20% in *Rag1^-/-^* and 10% in WT and *Tcrd^-/-^* of cells expressing IL-22 and IL-17A also co-express IL-13 (**Fig. 4E and F**).

## DISCUSSION

Experimentally combining scRNA-seq, ATAC-seq, and *in vivo* fate mapping in the psoriasis mouse model with new analytical approaches, we showed the presence of previously undescribed naïve/quiescent-like tissue-resident ILCs and the ability of activated ILC2s to differentiate to pathological ILC3s. We further discovered a novel subset of ILCs expressing IL-13 and IL-22/IL-17A in response to IL-23 stimulation. Our work highlights the limitation of experimental and computational analyses of immune cells that treat them as discrete immune “types”, when immune cells may share biological signals and span continuous spectra. In our system, we did not observe any discrete boundaries in single-cell expression profiles that neatly partitioned naïve/quiescent-like ILCs from activated type 2 cells, or type 2 cells from type 3 cells. Rather, the entire population of skin-resident ILCs was functionally reconfigured and its spectrum shifted by disease induction. Indeed, imposing stress on an immune cell population may allow rapid shifting of such a spectrum towards alternative cell fates (Tusi, Wolock et al. 2018), and pathways similar to those we uncovered in the skin may play roles in other tissues. Importantly, this also suggests that studies of ILCs sorted on expression of specific cytokines, such as IL-5 (Ricardo-Gonzalez, Van Dyken et al. 2018), may not have fully assessed this larger continuum. This model substantially revises previous interpretations and can provide a unified framework for some observations in other systems, such as “functional compartmentalization” within ILC types and gut ILCs that could not be readily assigned to a single ILC type (Gury-BenAri, Thaiss et al. 2016). These studies did not report a differentiation from ILC2 to ILC3, (but rather reported that a core ILC2 module was robust to antibiotic perturbation, albeit with increased expression of genes associated in homeostasis with ILC3s (Gury-BenAri, Thaiss et al. 2016), which may reflect tissue-specific differences in ILC features (Ricardo-Gonzalez, Van Dyken et al. 2018). Computational models and biological interpretations that allow for such fluidity, including topic modeling, are thus valuable for uncovering biological phenomena because they highlight signals, such as, in our case, type 2 activation, shared by cells in distinct clusters, and reveal drivers of heterogeneity among cells within a single group, such as the ILC “cloud”. This type of presentation is consistent with recent studies of HSCs, where individual precursors have probabilistic fate maps, tilted towards but not commited to specific outcomes (Carrelha, Meng et al. 2018, Laurenti and Gottgens 2018). Such approaches should be valuable in uncovering how tissue-resident ILCs, and other cell types, may globally respond to a stimulus, and undergo dynamic, plastic activation to reach the necessary state for shaping the tissue landscape.

## METHODS

### Mice

C57BL/6, *Tcrd*^-/-^ and Rosa26^flox-stop-floxYFP^ Ai3(RCL-EYFP) mice were purchased from the Jackson Laboratories. *Rag1^-/-^* and *Rag2^-/-^IL2rg^-/-^* were purchased from Taconic Biosciences. IL-5 Cre, dTomato (Red5/R5) from Dr Locksley laboratory. The IL-5 fate reporter in this work was generated by crossing Red5 with Ai3(RCL-EYFP) with IL-17A^GFP^ (Esplugues, Huber et al. 2011) and IL-22^BFP^ generated in our laboratory. In order to maximize the Cre recombination and increase the signal of Rosa26^YFP^ positive cells, we used homozygous IL-5^dTomato,Cre^. We observed little to no difference in IL-23 induced skin thicknening (**Fig. S4A**).

All mice were kept under specific pathogen-free (SPF) conditions in the animal facility at Yale University. Age- and sex-matched littermates between 10 to 14 weeks of age were used for all experiments. Unless with special instructions, mice were randomly assigned to different experimental groups and each cage contained animals of all different experimental groups. Both male and female mice were used in experiments. Animal procedures were approved by the Institutional Animal Care and Use Committee (IACUC) of Yale University. Preliminary experiments were tested to determine sample sizes, taking available recourses and ethical use into account.

### Psoriasis model

The psoriasis model used in this study is based on rIL-23 subcutaneous injections. The 500ng in 20μl of rIL-23 (provided by Abbvie or purchased from R&D Systems [scRNAseq experiments]) was injected daily into the ear skin of anesthetized mice in 4 consecutive days. As a control 20μl of PBS was used with the same injection intervals. For the second challenge experiment, we waited 10 days, monitoring the skin thickness before repeating 4-day injection regimen. Skin thickness was measured daily with calipers. When indicated, FTY720 (1mg/kg) was dissolved in PBS and administered i.p on day −1, 1 and 3 of the experiment. Skin tissue was collected on day 5 for histology imaging, flow cytometry analysis or cell sorting.

### Isolation of skin lymphocytes

Ventral and dorsal dermal sheets of ears were separated, minced and incubated in RPMI medium containing 0.4mg ml^-1^ Liberase TM (Roche Diagnostics) and 60ng/ul DNAseI (Sigma). After digestion, the suspension was passed through and further mechanically disrupted with syringe plunger and a 70uM cell strainer. Lymphocytes were enriched by gradient centrifugation in 27.5% Optiprep solution (Sigma) and RPMI medium containing 5% Fetal Bovine Serum. Spleens were mechanically disrupted using a syringe plunger in complete RPMI. Cells were filtered through 70-μm nylon mesh and washed

### Flow cytometry and cell sorting

Mouse ILCs were stained with monoclonal antibodies to CD45.2, CD90.2, lineage (CD4, CD8, CD11b, CD11c, CD19, B220, NK1.1, Ter119, Gr1, FcEr1a), TCRβ, TCRγ, CD3ɛ. For intracellular cytokine staining, cells were re-stimulated for 6 h at 37°C with phorbol 12-myristate 13-acetate (PMA) (Sigma, 50 ng ml^−1^) and ionomycin (Sigma, 1 μg ml^−1^) in the presence of Golgistop (BD Bioscience) added after initial 2h of stimulation. Next, cells were fixed and stained with BS Cytofix/Cytoperm reagent (BD Biosciences) according to the manufacturer’s protocol. Intracellular cytokines were stained with antibodies to IL-13, IL-17A and IL-22. Total ILCs were sorted as live, CD45+, CD90+, lin- (CD4, CD8, CD11b, CD11c, CD19, B220, NK1.1, Ter119, Gr1, FcEr1a), CD3*ε*- and TCR γ/δ - cells into PBS/0.2%FBS.

### *In-vitro* ILC cultures

For *in vitro* experiments, 5,000 ILCs were cultured per well of a 96-well round bottom plate in Click’s medium with 10 ng ml^-1^ IL-2 (R&D Systems) and 25 ng ml^-1^ IL-25 (R&D Systems) with 10 ng ml^-1^, IL-33 (R&D Systems) or IL-23 25 ng ml^-1^ (Provided by Abbvie) with TGFβ 10 ng ml-1 (R&D Systems) and IL-1β 10 ng ml^-1^ (R&D Systems). Cells were collected for RNA extraction and qRT-PCR after 5 days of culture in 37°C and 5%CO_2_.

### Adoptive ILC transfer

Total skin ILCs were FACS purified and collected to PBS 5% serum. Cells were washed twice with 1x PBS and injected (10,000 cells per mouse in 100ul) into retro-orbital vein of anesthetized *Rag2^-/-^IL2rg^-/-^* mice. IL-23 injection experiments were performed 14 days after the transfer.

### RNA extraction and Quantitative Real time PCR (qRT-PCR)

RNA from *in vitro* cultures was isolated with RNeasy Mini Kit (QIAGEN) and qPCR was performed using KAPA Probe Fast qPCR Master Mix 2x Kit (Kapa Biosystems, Wilmington, MA) with TaqMan probes (Applied Biosystems) in a StepOne cycler (Applied Biosystems, Carlsbad, CA). The CT values from duplicate qPCR reactions were extracted from the StepOne cycler (Applied Biosystems, Carlsbad, CA) onto Excel spreadsheets and were analysed with the relative quantification method 2^ΔΔCT^.

### ATAC-seq

Total ILCs sorted from naïve wild type mice were processed for ATAC-seq analysis according to previously published protocol (Buenrostro, Giresi et al. 2013) with the low cell number input version (~5,000 ILCs). Libraries from two independent experiments were sequenced on HiSeq2500 with 75bp paired end reads. Each sample was sequenced to a depth of 150 million reads.

### ATAC-Seq data analysis

Adapter sequences were trimmed using FASTX-Toolkit (version 0.0.13, http://hannonlab.cshl.edu/fastx_toolkit/), after which Bowtie2 (Langmead and Salzberg 2012) was used to align the reads to the mm10 genome. Picard tools (version 2.9.0, https://broadinstitute.github.io/picard/) were used to remove PCR duplicates. Bedtools was used to convert the bam file to a bed file, and all mapped reads were offset by +4 bp for the positive strand and −5 bp for the negative strand. Peaks were called for each sample using macs2 (Zhang, Liu et al. 2008) using parameters –nomodel –nolambda –shiftsize 75. ATAC-seq peaks were visualizaed with the Integrative Genomics Viewer (Robinson, Thorvaldsdottir et al. 2011, Thorvaldsdottir, Robinson et al. 2013) along with publicly available ChIP-seq via Cistrome DB (Liu, Ortiz et al. 2011).

### Single cell RNA-Seq

Sorted cells were washed with PBS/0.04% BSA and processed for droplet-based 3′ end massively parallel scRNA-seq: sorted ILCs were encapsulated into droplets, and libraries were prepared using Chromium Single Cell 3′ Reagent Kits v2 according to the manufacturer’s protocol (10X Genomics). scRNA-seq libraries were sequenced using a 75 cycle Nextseq 500 high output V2 kit.

### Single cell RNA-Seq data analysis

#### Initial data processing and QC

Gene counts were obtained by aligning reads to the mm10 genome using CellRanger software (v1.3) (10x Genomics).

To remove doublets and poor-quality cells, cells were excluded from subsequent analysis if they were outliers in their sample of origin in terms of number of genes or number of unique molecular identifiers (UMIs), which eliminated 5.8–7.9% of cells per sample (**Fig. S4B**), or outliers across all samples in percentage of mitochondrial genes, which eliminated at most 0.5% of remaining cells (**Fig. S4C**). Sample-specific cut-offs ranged from 575–2,400 genes per cell for the *Rag1*-/- untreated sample to 850–3,100 genes per cell for the WT induced sample.

#### Normalization

To normalize gene counts, we used a scaling factor that reflected the expected number of UMIs in each sample (**Fig. S4D**), rather than scaling all cells to a constant size, as in TPM (Wallrapp, Riesenfeld et al. 2017) Let *w_s_* be the mean number of UMIs per cell in sample *s*. UMI counts for cells in sample *s* were scaled to: 10, 000 × (*w_s_*/*w*WT _naive_)

Taking the log of scaled UMI counts gives the normalized expression values referred to as logTPX.

#### Determination of variable genes

We fit the count data to a null model based on a negative binomial distribution that explains the expected technical variation for each gene, given its expression level, as previously described(Pandey, Shekhar et al. 2018). A gene was considered to exhibit non-technical variability if it had mean counts above 0.005 and a coefficient of variation at least log(0.5) times that predicted by the null model (**Fig. S4E**). We performed variable gene selection separately for each sample as well as for pooled samples from WT mice and, separately, from *Rag1*^-/-^ mice. To reduce downstream technical effects of the variation in extremely highly expressing genes, we then removed any genes that had mean counts above 4 in WT or, separately, *Rag1*^-/-^ cells (these were mostly ribosomal protein genes). The resulting conservative set of 271 genes was then used for the singular value decomposition (SVD). We chose this approach to ensure that noisy variable gene selection was not a cause of the heterogeneity in the “cloud”. Note that downstream results were qualitatively similar and robust to several parameter settings, which yield variable gene sets of very different sizes, as well as to other selection approaches (including the FindVariableGenes() function in Seurat) (Butler, Hoffman et al. 2018).

#### Dimensionality reduction, clustering, and visualization

We computed an SVD on *z*-scored variable genes, as determined above, using Seurat’s RunPCA() function, with the “weight.by.var” parameter set to FALSE (Butler, Hoffman et al. 2018). Assessing the decrease in marginal proportion of variance explained with larger components, we selected the top 18 eigenvectors for subsequent analysis, and confirmed that the resulting analyses were not sensitive to this exact choice. We used these components with Seurat’s FindClusters() and RunTSNE() functions, with other parameter settings set to default, to cluster the cells, and to separately create a *t*-stochastic neighborhood embedding (tSNE) for visualization, respectively. As previously described, FindClusters() optimizes a modularity function on a *k*-nearest-neighbor graph computed from the top eigenvectors.

#### Removal of non-ILC clusters

Based on expression of marker genes across clusters, we determined that a few very distinct clusters were unlikely to be ILCs: cells in those clusters had little expression of *Ptprc* (CD45), and high expression of *Col1a2*, or *Tie1* and *Pecam1,* or *Krt15.* Cells from these non-ILC clusters were removed, and the steps of normalizing the data, selecting variable genes, performing PCA, and creating a tSNE were repeated as before, but the top 20 components of the SVD were used for subsequent analysis. After these steps, 18,852 cell profiles remained, with 4,619–4,857 cells per sample.

#### Topic modeling

We fit an LDA topic model on the full, sparse counts matrix (18,852 cells and 27,998 genes) using the FitGoM() function from the CountClust R package (Dey, Hsiao et al. 2017), with the number of clusters *K* set to 15 and the “tol” tolerañce parameter set to 10. This package is heavily based on the maptpx R package, which implements a posterior maximization approach to fitting the model (Taddy 2012). Some approaches to selecting an appropriate value of *K* rely on having labeled training data for the model. Since we do not have such a model, we fit the model for a range of values and computed the Akaike and Bayesian information criteria (AIC and BIC) using the estimated likelihood returned by FitGoM() (**Fig. S2A**). Since AIC and BIC risk under- and over-penalizing the fit, respectively, we selected a value of *K* at a point where the AIC curve had begun to decrease less steeply and the BIC curve had begun to climb.

#### Diffusion maps

To select cells and genes for the construction of diffusion maps, a cell was considered “highly weighted” for a topic if its weight for the topic was above a topic-specific threshold capturing the upper tail of the distribution (**Fig. S3A and B**). The analysis is not sensitive to the exact choice of threshold. Cells were used in the large diffusion map (**Fig. 3A**) if they were highly weighted for any of topics 2, 4, 8, 11, 13, or 15, but not 6 or 7 (**Fig. S3B and C**). A gene was considered to be in the “top *n* genes” for a topic if it was returned by the CountClust function ExtractTopFeatures(), which selects genes that are most critical for separating one topic from the others (similar to differential expression analysis between clusters), with the following parameter settings: top_features=*n,* method=“poisson”, options=“min”, shared=TRUE. For visualization, the “Score” shown for top genes (**Fig. 2C, Fig. S2D**) was computed as 100\**x,* where *x* is the Kullback-Leibler divergence score output by ExtractTopFeatures(), and then plotted on a logarithmic scale. Genes were included in the large diffusion map if they were in the top 50 genes for topics 2, 4, 8, 11, 13, or 15, but not in the top 5 genes for any other topics. For the smaller diffusion map (**Fig. 3C**), cells and genes were selected in an analogous way, but only for the three topics 8, 13, and 15. Overall, the larger diffusion map was computed on 7,888 cells and 245 genes, and the smaller one on 3,785 cells and 130 genes. To build the diffusion map, we gave the expression data for these cells and genes as input to the DiffusionMap() function from the destiny R package (Angerer, Haghverdi et al. 2016), with parameter settings *k*=50 and sigma=“local”.

## Acknowledgments

We would like to thank Ania Hupalowska and Leslie Gaffney for help in manuscript preparation. This work was supported in part by grants (YAP-013-2015) provided by AbbVie (RAF, PB). Work supported by the Klarman Cell Observatory (AR) and HHMI (AR and RAF). AR is an SAB member of ThermoFisher Scientific, Syros Pharmaceuticals, and Driver Group and a founder of Celsius Therapeutics.

---

**Figure S1.**
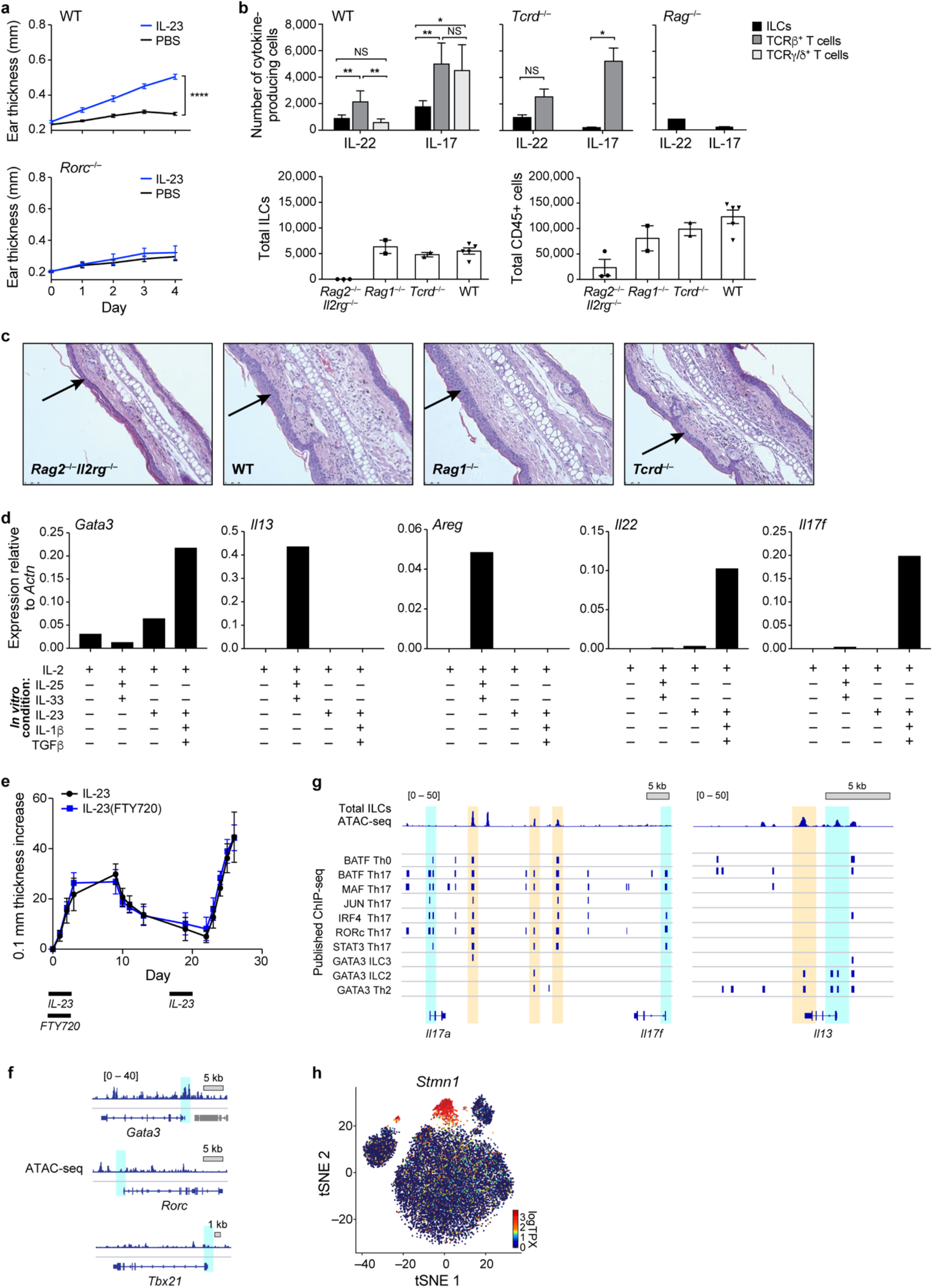
Characterization of skin immune cells to IL-23 induction, Related to Figure 1. (A) Increase in ear skin thickness is significantly higher in response to IL-23 treatment than PBS vehicle and is dependent on *Rorc*. Increase in ear thickness (y axis, mm) following treatment with IL-23 (blue) or PBS vehicle (black) in WT (top) or *Rorc^-/-^* (bottom) mice. (B and C) Immune cell composition and skin phenotype in different mouse genotypes. (B) Top: Number of cells (y axis) producing IL-22 or IL-17 (x axis) among ILCs (black bars), γδT cells (grey bars) and γδ T cells (white bars) in WT, *Tcrd*^-/-^ (lack γδ T cells), *Rag1*^-/-^ (lack all T and B cells), and *Rag2*^-/-^ *Il2rg*^-/-^ mice (also lack ILCs) mice. Bottom: Number of total CD45+ (left, y axis) or total ILCs (right, y axis) in WT, *Tcrd*^-/-^, *Rag1*-/-, and *Rag2*^-/-^ *Il2rg*^-/-^ mice (x axis). (C) H&E stains of ear sections in each genotype except *Rag2*^-/-^ *Il2rg*^-/-^ mice. Arrows: Acanthosis. (D) Expression of type 2 and type 3 related genes in cultured naïve skin ILCs. Shown are relative expression levels (y axis, by qPCR) in ILCs cultured with different cytokines (x axis, table at bottom). (E) FTY720 treatment does not impact increased susceptibility to a secondary challenge with IL-23. Skin thickness (y axis, 0.1mm) over time (x axis, days) in mice following treatment with either IL-23 or IL-23 and FTY720. Bottom bars: period of primary (left bar) and secondary (right bar) challenge. (F and G) ATAC-seq of sorted skin ILCs from untreated mice. Mapped ATAC-seq reads from sorted skin ILCs from untreated mice (top tracks) at different loci (bottom tracks) of interest. Blue shaded areas: TSS; Beige shared areas: open chromatin peaks at key TF binding sites, previously identified in CD4+ T cell ChIP-seq data (middle tracks). (F) Open chromatin peaks at TSS of *Gata3* (associated with mature ILC2) but not *Rorc* (ILC3) and *Tbx21* (ILC1). (G) Open chromatin peaks at TF binding sites in the *Il17a and Il17f* promoter (beige), and at the TSS of *Il13* but not *Il17a,Il17f* (blue). (H), Cluster B reflects cycling ILCs. tSNE of 27,998 single ILC profiles (dots) colored by expression level (logTPX, color bar) of *Stmn1*.

**Figure S2.**
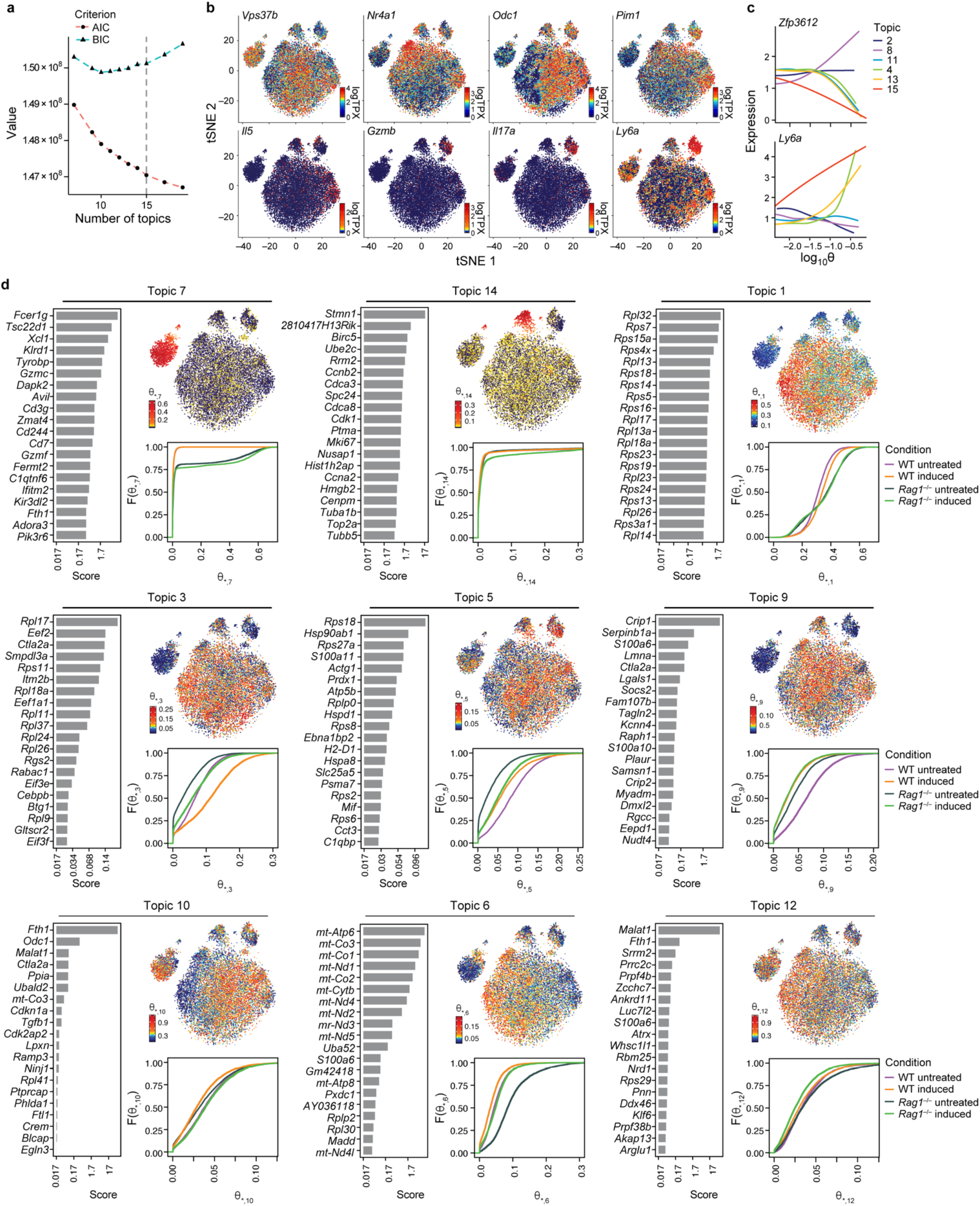
Topic modeling also distinguishes cluster-specific, cell size, and cell quality related topics, Related to Figure 2. (A) Selecting the number of topics. Akaike Information Criterion (AIC, red) and Bayesian Information Criterion (BIC, blue) value (y axis) for a range of the number *K* of topics. *K=*15 (dotted line) is at a point where the AIC curve decreases less steeply and the BIC curve begins to rise. (B and C) Expression of example genes associated with key topics. (B) tSNE of 27,998 single ILC profiles (dots) colored by expression level (color bar, logTPX (**Methods**)) for genes in Topics 2, 4, 8, 11, 13, and 15. (C) Gene expression (y axis, logTPX) as a function of the topic weight (log θ_*_,*j*, x axis), for each of these topics (color), for repressive gene *Zfp36l2* and activation-associated gene *Ly6a*. (D) Summary of remaining topics not included in Fig. 2c. For each topic shown are a bar plot of top scoring genes (y axis), ranked by a score (x axis, logarithmic scale) of how well the gene distinguishes this from other topics (**Methods**); a tSNE (as in B) with cells colored by the topic’s weight in the cells (column *j* of the cell-by-topic weight matrix θ (θ_*_,*j*) for Topic *j*); and a graph of the empirical cumulative density function (y axis) of topic weights θ_*_,*j* (x axis) for cells grouped by treatment or genotype (as in Fig. 2c).

**Figure S3.**
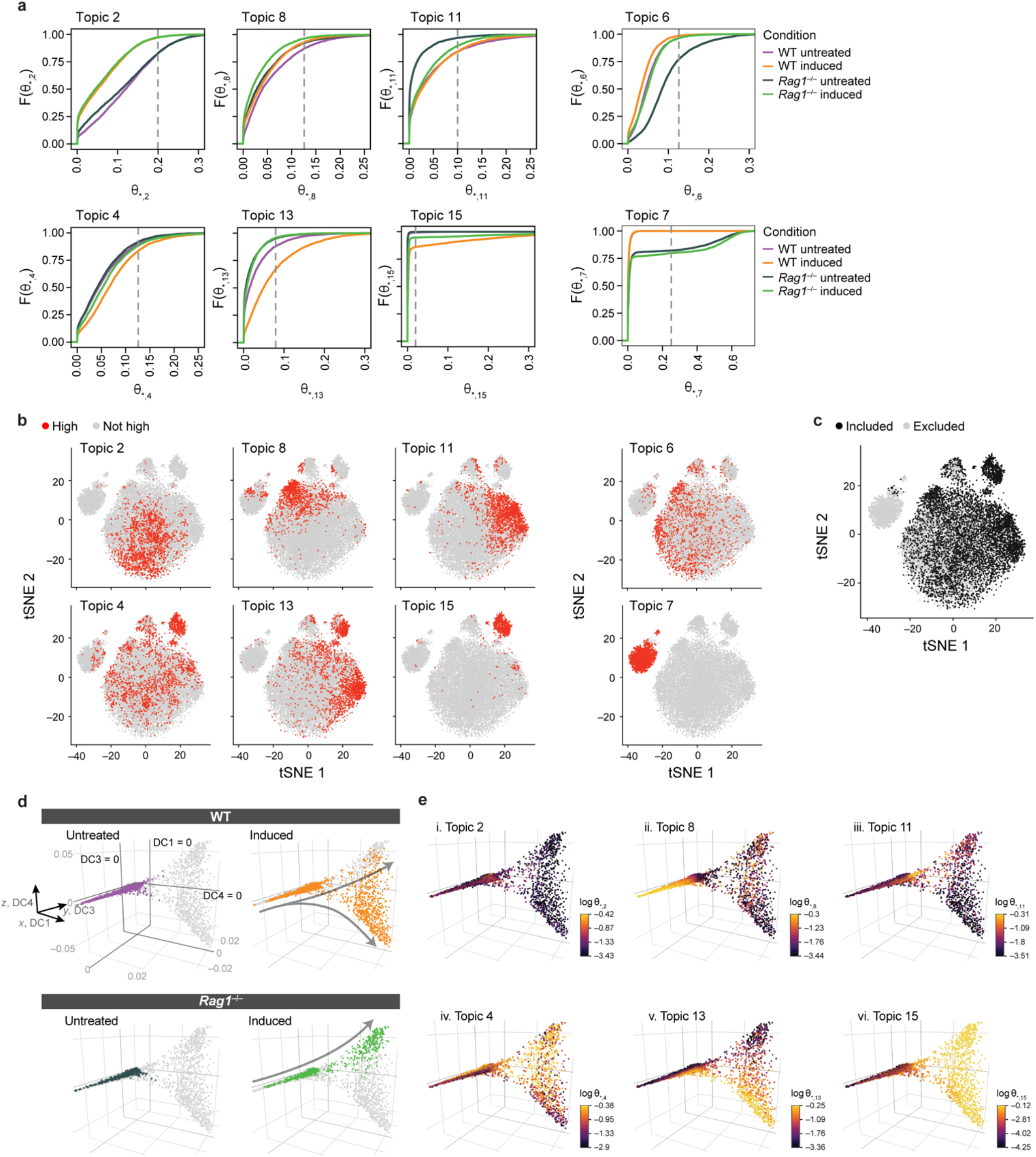
Diffusion map analysis based on topic model highlights an IL-23–induced dynamic trajectory, Related to Figure 3. (A-C) Cell selection for diffusion map in Fig 3A and B. (A) Chosen topic weight thresholds. Empirical cumulative density function (y axis) of topic weights θ_*_,*j* (x axis) of cells grouped and colored by *in vivo* treatment and genotype. Dotted line: topic weight threshold. (B and C) Cells with high weights in at least one key topic are chosen for the diffusion map. tSNE of 27,998 single ILC profiles (dots), with cells colored if they are weighted above the corresponding topic threshold from A (B, red) and chosen for the diffusion map (C, black), if they are highly weighted for Topics 2, 8, 11, 4, 13, or 15, but not for Topics 6 or 7. (D and E) Topics 8, 13, and 15 highlight a potential naïve-induced trajectory across DC1. Plots of DC1 (x axis), DC3 (y axis), and DC4 (z axis) show cells (dots) colored by either *in vivo* treatment and genotype (D) or by topic weight (log θ_*_,*j*, color bar) (E). Gray arrows (D) indicate an implicit direction of induction.

**Figure S4.**
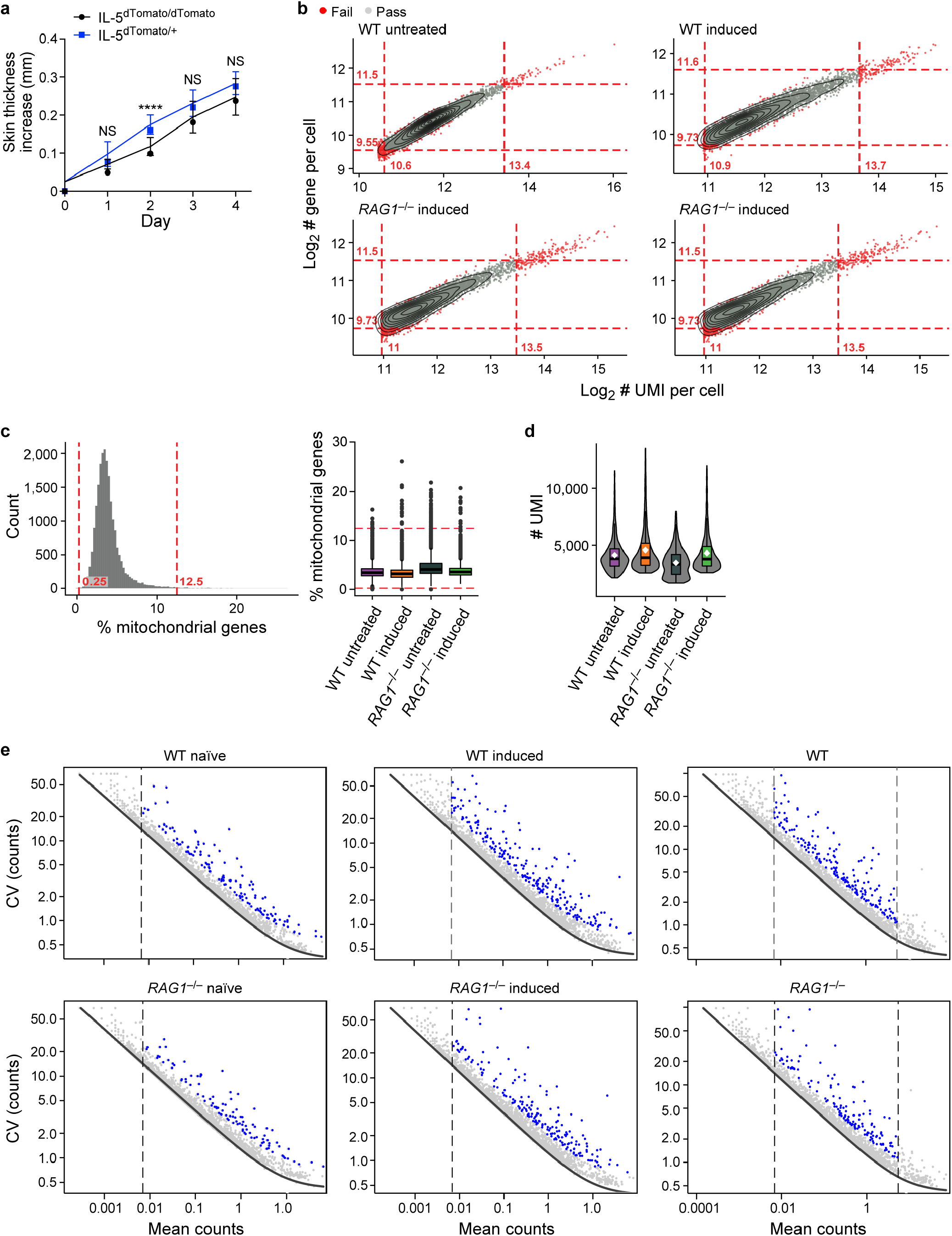
Computational and experimental quality control and data processing. (A) IL-23 skin injection model in *Il5*^dTomatoCre^ (Red5) mouse strain. Increase in skin thickness (mm, y axis) over time (days, x axis) in homozygote Red5/Red5 mouse strain lacking expression of IL-5 cytokine (black) and Red5/+ mouse (blue) shown little difference. (B and C) Quality control and filters in scRNA-Seq. (B) Minimum and maximum thresholds (red) of log UMI counts (x axis) and log gene counts (y axis) for each condition. Cells (dots) that were filtered out are marked in red. (C) Left: Histogram of the % of mitochondrial genes detected across all cells. Right: Box plot of the % of mitochondrial genes of all detected genes in each sample type. Dashed lines: thresholds. (D) Distributions for the number of UMI counts in each sample, used to compute logTPX (**Methods**). (E) Variable gene selection. For each sample (panel) shown are the coefficient of variation (CV, y axis) as a function of mean counts (x axis) for each gene (dot). Black curve: null model (**Methods**). Blue: genes with sufficiently greater CV than in the null model, which are retained as variable genes.

